# Beyond demographic buffering: Context dependence in demographic strategies across animals

**DOI:** 10.1101/2021.06.08.447594

**Authors:** Omar Lenzi, Arpat Ozgul, Roberto Salguero-Gómez, Maria Paniw

**Author notes:** Corresponding author’s.

## Abstract

Temporal variation in vital rates (*e.g*., survival, reproduction) can decrease the long-term mean performance of a population. Species are therefore expected to evolve demographic strategies that counteract the negative effects of vital rate variation on the population growth rate. One key strategy, demographic buffering, is reflected in a low temporal variation in vital rates critical to population dynamics. However, comparative studies in plants have found little evidence for demographic buffering, and little is known about the prevalence of buffering in animal populations. Here, we used vital rate estimates from 31 natural populations of 29 animal species to assess the prevalence of demographic buffering. We modeled the degree of demographic buffering using a standard measure of correlation between the standard deviation of vital rates and the sensitivity of the population growth rate to changes in such vital rates across populations. We also accounted for the effects of life-history traits, *i.e*., age at first reproduction and spread of reproduction across the life cycle, on these correlation measures. We found no strong or consistent evidence of demographic buffering across the study populations. Instead, key vital rates could vary substantially depending on the specific environmental context populations experience. We suggest that it is time to look beyond concepts of demographic buffering when studying natural populations towards a stronger focus on the environmental context-dependence of vital-rate variation.

## Introduction

Individuals in natural populations can experience large variation in vital rates (*e.g*., survival, growth, or reproduction), which implies that they experience, and are affected by, different environments over their lifetime (Boyce et al. 2006). Such a variation has been shown to negatively affect population performance (Gillespie 1977, Tuljapurkar 1982, Moore and Huntington 2008, Regehr et al. 2010), increasing the probability of extinction (Lewontin and Cohen 1969, Drake and Lodge 2004, Adler and Drake 2008). To avoid the negative consequences of such variation, species have evolved demographic strategies, i.e., differences in the temporal patterns of vital rates. Studying these strategies has been important to support wildlife management and conservation actions (Caswell 2007, Lawson et al. 2015, McDonald et al. 2017).

Following the *demographic buffering hypothesis* (sometimes also referred to as canalization [*sensu* Gaillard and Yoccoz 2003]), it is expected that vital rates contributing most to population growth rate (λ) should be buffered the most against environmental variation, while vital rates that contribute little to population dynamics are expected to vary more freely because they are not as strong a target of selection (Fig. 1) (Pfister 1998, Hilde et al. 2020). A key assumption of this hypothesis is that an increase in vital-rate variation decreases λ *(Morris et al. 2008, Koons et al. 2009)*. The demographic buffering hypothesis has been tested on and found support among some plants (Pfister 1998, Morris and Doak 2004, Burns et al. 2010, Li and Ramula 2015) and a few animals, such as *Falco peregrinus* (peregrine falcon), *Leptonychotes weddellii* (Weddell seal), and *Rangifer tarandus platyrhynchus* (Svalbard reindeer) (Pfister 1998, Rotella et al. 2012, Bjørkvoll et al. 2016).

**Figure 1.**
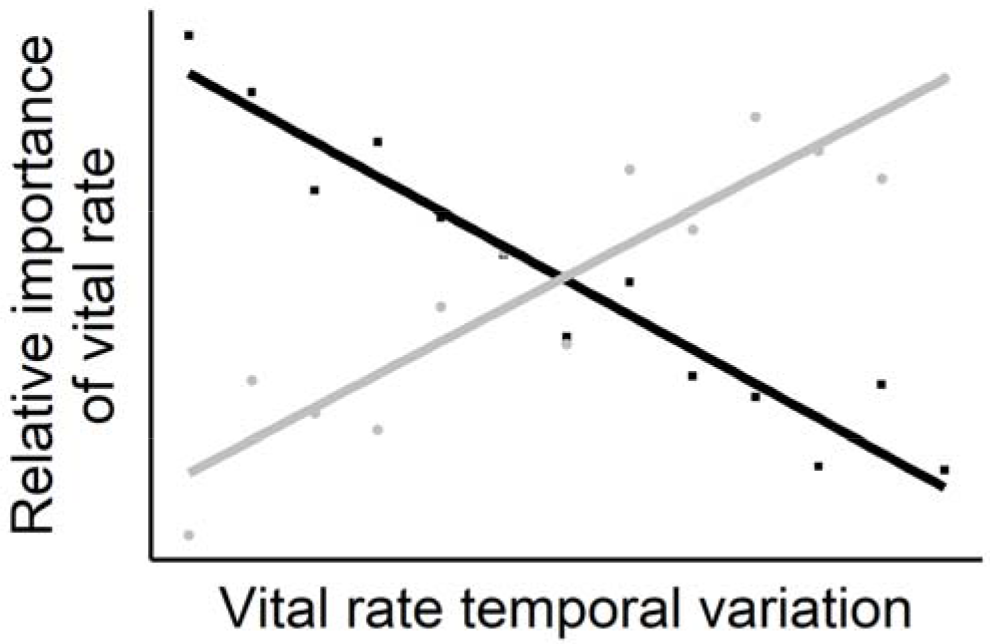
Graphical representation of the demographic buffering (black) and the demographic lability hypotheses (gray). The demographic buffering hypothesis predicts lower temporal variation in the vital rates that contribute the most to the population growth rate, whereas the demographic lability hypothesis predicts that the most important vital rates track environmental changes, therefore showing the most temporal variation. Each square or circle represents a vital rate, for which its variation over time, as well as its relative importance for the population growth rate, are calculated.

Comparative studies in plants have recently cast doubt on the universality of demographic buffering. It has been suggested that evidence for this demographic strategy may often be due to statistical artefact arising when calculating vital-rate variation rather than due to biological mechanisms (Burns et al. 2010, Bjørkvoll et al. 2016, McDonald et al. 2017). In addition, selection can favor phenotypic plasticity in response to environmental fluctuations, whenever a nonlinear relationship between a vital rate and the environment enhances the temporal mean of the vital rate leading to an increased population growth rate (Koons et al. 2009) (Fig. 1). Under such circumstances, the *demographic lability hypothesis* suggests that the average population fitness increases when variation in the most important vital rates tracks environmental variation (Koons et al. 2009, Jongejans et al. 2010) and implies a positive correlation between the contribution of vital rates to population dynamics and their variation (Drake 2005, McDonald et al. 2017). Still, while providing important evidence of how species may respond to environmental change, the relationship between vital-rate variation and vital-rate contribution to average population fitness has not been explored systematically in animals.

Here, we examine the prevalence of demographic buffering across animal populations. Based on previous studies on plant populations (Burns et al. 2010, McDonald et al. 2017), we expected variable demographic strategies in animals too, where only some populations would show demographic buffering. As has been suggested in previous studies (Morris et al. 2008, McDonald et al. 2017), we expected that those populations showing buffering would be long-lived animals, while short-lived animals where reproduction plays a more important role in the life cycle and species that track environmental change would be least likely to be buffered (Ripley and Caswell 2006, Koons et al. 2009). Moreover, we expected that highly iteroparous organisms would be more likely to display demographic buffering because one important way in which organisms can buffer environmental variation is by spreading reproduction across more reproductive events, with fewer but more successful offspring (Miller et al. 2011). To test our expectations, we examined 31 natural populations from 29 animal species worldwide (Table S1). For each population, we assessed the relationship between standardized vital-rate variation and vital-rate contribution to population fitness using Bayesian mixed-effects models. In the models, we accounted for two key life-history traits: (i) age at first reproduction and (ii) spread of reproduction, which determine two independent axes of life-history strategies in animals and plants (Paniw et al. 2018, Healy et al. 2019).

## Methods

### Study populations

To obtain demographic information with temporally replicated vital rates, we accessed the COMADRE Animal Matrix Database version 2.0.1 (http://www.compadre-db.org) (Salguero-Gómez et al. 2016), as well as other supplemented peer-reviewed publications (Table S1). We chose matrix population models (MPMs) (i) not subject to experimental manipulations, as we were interested in natural temporal variation; (ii) that represented at least three annual transitions in order to calculate variation in vital rates; (iii) that were built from a single wild population; and (iv) that had entries for reproduction, with no missing values. Our selection criteria resulted in 27 populations from 25 animal species (Table S1). In addition, we obtained vital rates from a natural population of snow voles (*Chionomys nivalis*) (Bonnet et al. 2017), a population of meerkats (*Suricata suricatta*) (Ozgul et al. 2014), a population of yellow-bellied marmots (*Marmota flaviventer*) (Paniw et al. 2020) and a population of woodland caribou (*Rangifer tarandus caribou*) (DeCesare et al. 2012) (see Data S1).

### Standardizing sensitivities and variances

To determine the prevalence of demographic buffering in each study population, we determined the relationship between the variation in a vital rate of interest and its influence on population growth rate (*i.e*., sensitivity of the growth rate to changes in a vital rate) (de Kroon et al. 1986, Caswell 2001). To calculate vital-rate variation and sensitivities, we adapted a method developed by McDonald et al. (2017) (see `corrected sensitivities and variances.R` in Supporting Materials). Briefly, this method is based on the calculation of corrected standard deviations and scaled sensitivities, thereby accounting for the variance constraints that arise in vital rates. For instance, in vital rates constrained between zero and one, such as survival and stage transition probabilities, the variance depends on the mean value and is therefore only comparable with vital rates that do not have an upper limit, such as reproduction, if a scaling is applied (Link and Doherty 2002).

The first step in the calculation of the corrected standard deviations was the extraction of the underlying vital rates from MPM elements, which can comprise several vital rates (*i.e*., survival and stage transition probabilities) (Morris and Doak 2004). We calculated stage-specific survival as the column-sum of all the non-reproduction elements in the MPM. Conditional on survival, we then extracted progression between stages *j* and *i* from time *t* to *t*+1. We note that retrogression (Franco and Silvertown 2004) was not present in any of the MPMs used in this study. Reproduction was calculated as the mean offspring number produced by an individual in a given class. The second step was to calculate the sensitivities (S) of population growth rate (λ) to changes in vital rates (*vr*). We first built the average (across year) MPM for each population and then calculated the vital rate-specific sensitivities following methods detailed by Caswell (2001) and Franco and Silvertown (2004):

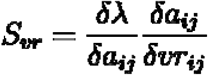

where *a* is the matrix element in the *i,j^th^* position in an MPM. Sensitivities assume independence of vital rates, *i.e*., a change in a given vital rate would affect λ independently of other vital rates. However, vital rates covary, and this covariation can alter our interpretation of the effects of vital-rate variation on population dynamics (Doak et al. 2005). To account for covariation in vital rates, we additionally calculated integrated sensitivities as proposed by van Tienderen (1995). Integrated sensitivities represent the sum of the direct effect of a vital rate on λ and its indirect effect via correlations with other vital rates (for details see `corrected integrated sensitivities and variances`.R in Supporting Materials).

To account for the constraints of vital-rate variances, we calculated corrected standard deviations as the standard deviation on the logit of each binomial vital rate (survival and stage transition probability) and on the log of zero-constrained vital rates (reproduction). Similarly, we scaled vital-rate sensitivities by calculating the variance-stabilised sensitivity (VSS) according to McDonald et al. (2017), adapted from Link and Doherty (2002):

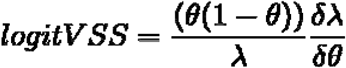

where λ is the dominant eigenvalue from the average (across year) MPM, and *θ* are the binomial vital rates. For reproduction, where the variance is only bounded downwards by zero, the VSS correction corresponded to calculating the elasticities of the population growth rate to changes in these vital rates (McDonald et al. 2017) (Data S1). We inspected visually whether associations between the variance and mean of vital rates were present in our data and whether link-scaling removed these within-species associations (Fig. S1).

### Demographic strategies

After standardizing vital-rate variances and population-growth sensitivities, we evaluated the degree of buffering among the study populations in two steps. First, we calculated, for each study population, the Spearman correlation coefficient between the corrected standard deviations and sensitivities, including integrated sensitivities (Supporting Materials). As a measure of the strength of the association, we calculated 95% confidence intervals (C.I.) of the correlation coefficients, using the function *spearman.ci* from the package *RVAideMemoire* (Hervé 2019). However, this procedure could only be done for the 19 populations with five or more vital rates (Figs. S2 & S3), since the estimated C.I. are not reliable with fewer data points.

Next, to better account for the uncertainty in correlation estimates while integrating species with few vital rate estimates, we fitted a generalized mixed effect model with log-transformed standardized sensitivities as a response and standardized vital-rate variations as predictor. We included a random species effect on the slope of the model, the latter providing a measure of demographic strategy (buffering if negative and lability if positive) akin to the Pearson’s correlation coefficients. To test for a phylogenetic effect, we associated the species with a phylogenetic tree (Hadfield 2010). We obtained this tree using the Open Tree of Life (Hinchliff et al. 2015) with the *rotl* R package (Michonneau et al. 2016) as well as the *phytools* R package (Revell 2012). For the one species where we modelled three different populations (*Macropus eugenii*), we added each population as a separate branch to the tree (see Fig. S4).

We tested whether the slope describing the relationship between vital-rate variation and population-growth sensitivities was affected by two life-history traits that describe independent axes of life-history strategies (Healy et al. 2019):

- Age at sexual maturity (*L*_□_): Number of years that it takes an average individual in the population to become sexually reproductive.
- Spread of reproduction across age classes (G): Gini index applied on the life table decomposition of MPMs; G□=□1 describes fully semelparous populations, where all individuals reproduce at the same age; G□=□0 describes an extremely iteroparous population, where all ages classes reproduce.

We accounted for the effect of phylogeny and adult body mass (Myhrvold et al. 2015) on both traits using a phylogenetically corrected mixed effect model following Healy and co-authors (2019), and incorporated the residuals from this model as an interaction term in the above-described demographic-strategy model. Details on the calculations can be found in the Supporting Materials (see Fig. S5). All modelling was performed using the *MCMCglmm* R package (Hadfield 2010). We used default priors as suggested in Healy et al. (2019) and ran three chains (three different runs of the *MCMCglmm* function) for a total of 1,100,000 iterations, sampling every 500th posterior value after a burnin of 100,000 iteration. We inspected the convergence of chains visually using trace plots and analytically by calculating the Gelman-Rubin diagnostic (Brooks and Gelman 1998).

## Results

There was little consistency regarding the prevalence of demographic buffering across the 31 studied populations. The relationship between the significance of a vital rate to population growth rate and its temporal variation differed starkly among populations (Fig. 2, Figs. S2 & S3). For most species, we could not discern a strong pattern of decrease in variability with increases in vital-rate significance (suggesting demographic buffering) or *vice versa* (suggesting demographic lability) (Fig. 2). Although in 23 of the 31 study populations the Spearman correlation coefficient between the significance of a vital rate to population growth rate and its temporal variation was negative, the 95 % C.I. indicated that this evidence for buffering was only substantial (95 % C.I. did not cross 0) for two species: *Macaca mulatta* and *Marmota flaviventer* (Fig. S2). For the remaining species, confidence intervals around mean estimates of correlation coefficients indicated that the demographic strategy could not be unequivocally determined. These results did not change when comparing vital-rate variation to integrated sensitivities where the covariation among vital rates was accounted for (Fig. S3).

**Figure 2.**
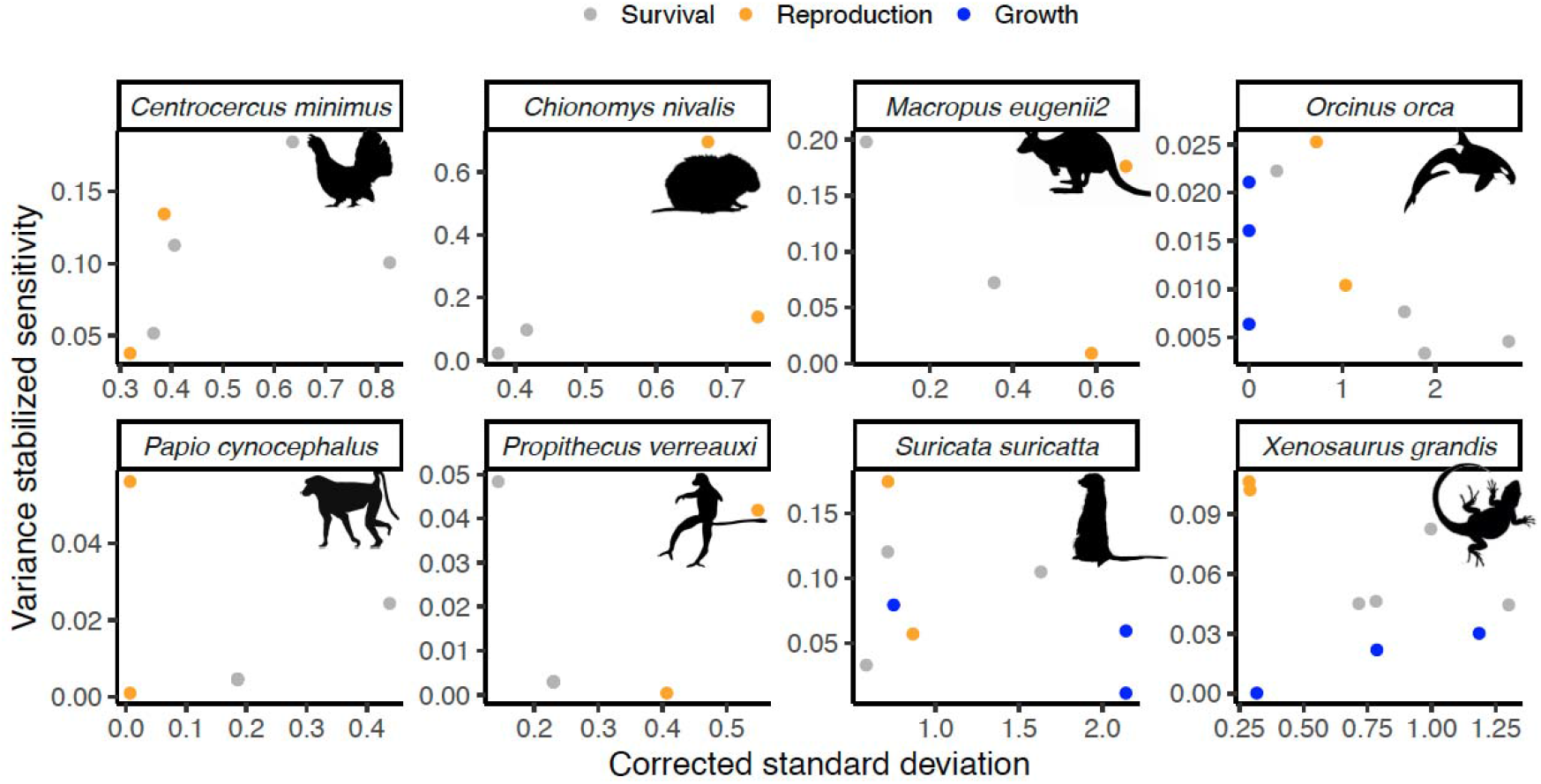
Temporal variation of stage-specific vital rates (*i.e*. the corrected standard deviation) and their importance to the population growth rate (*i.e*. the variance stabilized sensitivity) for eight study populations representing different families in the phylogenetic tree (for equivalent plots of all populations, see Fig. S2 and Fig. S3). We note that for the populations shown here, the demographic strategy could not be determined unequivocally (*i.e*., 95 % C.I. of Spearman correlation coefficient crossed 0).

The lack of consistent relationships between the significance of a vital rate to population growth rate and its temporal variation was confirmed by the results of our phylogenetically corrected mixed-effect models (Fig. 3). Both the phylogenetic effect (Fig. 3a) and species differences (Fig 3b) on the slope modelling variance stabilized sensitivities as a function of corrected standard deviations were negligible. Contrary to our expectation this slope also differed little as a function of age at sexual maturity (*L*_□_) and spread of reproduction (*G*). These results did not change when modelling integrated sensitivities as response (Fig. S5).

**Figure 3.**
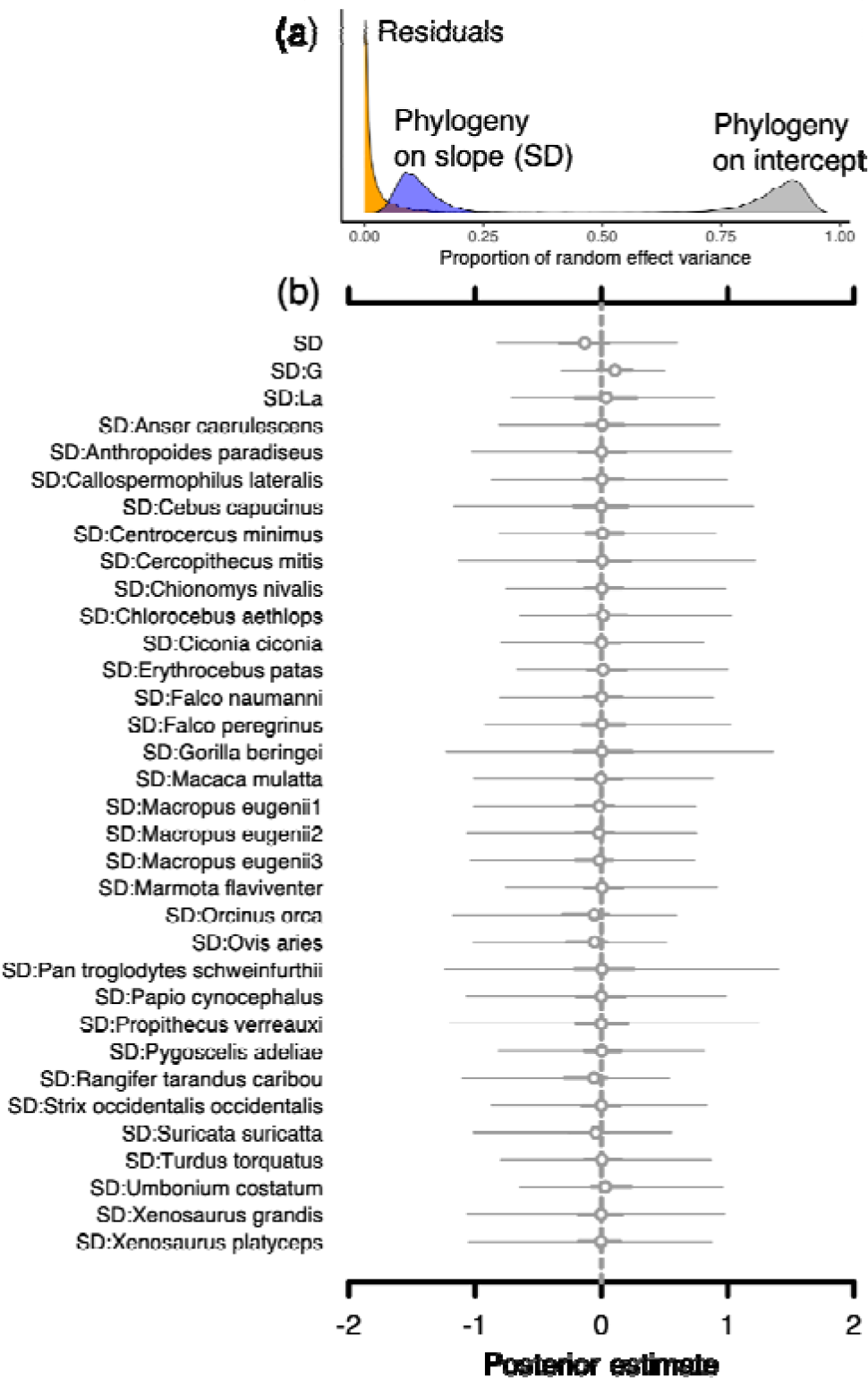
Effect of phylogeny (posterior parameter distribution) on model coefficients from the Bayesian mixed effect model describing the log(variance stabilized sensitivity of population growth) as a function of corrected vital rate standard deviations (SD) (a). Caterpillar plots (b) show the estimated fixed effects of SD, age at first reproduction (*L*_□_) and reproduction spread (*G*), while accounting for a phylogenetically corrected species effect on SD. Interactions among predictors are depicted by “:”. Points represent posterior medians. Parameters where 50% credible intervals (C.I.) overlap 0 are indicated by open circles. Thick lines represent 50% C.I.; thin lines represent 95% credible intervals.

## Discussion

In light of an impending extinction crisis, worsened by increasing environmental variation due to anthropogenic actions (Intergovernmental Panel on Climate Change 2014), an understanding of demographic strategies among natural populations to deal with such variation is important to guide future global-change research (Parmesan and Yohe 2003, Lawson et al. 2015). The assumption that all organisms show little variation in, or buffer, vital rates that are critical for population fitness has recently been questioned for plants (McDonald et al. 2017). Here, we show that such assumptions must be considered with more nuance for animals as well. Using the most common methods to assess buffering based on vital-rate variance and growth-rate sensitivity correlations, we found weak to no evidence for buffering in most examined natural populations, regardless of whether we considered standard or integrated sensitivities when calculating demographic strategies. The fact that accounting for covariation did not change our results is in line with previous studies that demonstrated little effect of accounting for vital-rate covariation on population dynamics (Morris et al. 2011, Compagnoni et al. 2016).

Large confidence intervals around estimates of correlation coefficients and mixed-effect model parameters demonstrate that the demographic strategy, regardless of the method used, could not be determined unequivocally (with the exception of two species that showed buffering). These large confidence intervals resulted -at least partly- from the limited number of vital rates available per population in matrix population models (Fig. S2), increasing uncertainties in parameter estimates. At the same time, even for species with > 5 vital rates, sensitivity-variance relationships varied greatly, with no clear pattern of the vital-rate variance decreasing with the “importance” of vital rates to population growth (suggesting buffering) or vice versa (suggesting lability). In many populations, most vital rates showed little variation regardless of their contribution to population growth, and the pattern of high variation and low sensitivity (or vice versa) was driven by one or two vital rates (Fig. S2). Our results on animals shadow those of McDonald et al. (2017) on plants, who also found little robust evidence for strong buffering, and suggest that sensitivity-variance relationships in vital rates are likely driven by population context and are challenging to address using current methods.

Our results show the limitations of generalizing demographic strategies (as we do here) without looking at the specific ecological context determining these strategies. For instance, various long-lived primates have previously been described as showing little sensitivity to environmental variation (Morris et al. 2011, Campos et al. 2017). However, once corrected for variance constraints, our results show that survival rates that contribute most to population growth rate may vary as much as or more than rates that contribute less to average population fitness. A higher vital-rate variation than would be expected might be caused by factors such as outbreaks of diseases, as was the case for the studied chimpanzee populations (*Pan troglodytes schweinfurthii*) (Pusey et al. 2008, Williams et al. 2008). However, vital-rate variation in primates was linked explicitly to climatic variation to show weak climate effects on critical vital rates (Campos et al. 2017), emphasizing the importance to consider explicit environmental drivers when assessing demographic strategies (Rodríguez□Caro et al. 2021).

Understanding the environmental context of vital-rate variation is not only important for long-lived species but also for ones that would be expected to show stronger lability given their short life span and relatively high reproductive output. For instance, both populations of the two *Xenosaurus* species might show little variation in key vital rates because they are territorial, leaving only rarely the microhabitats they occupy (Zúñiga-Vega et al. 2007). Such a strong territoriality, especially if microhabitat conditions do not vary strongly, might create a predictable and stable environment, where selection for tracking environmental change, *i.e*., lability, is not favored (Chevin et al. 2010, McDonald et al. 2017). As another example, meerkats might show little variation in important vital rates due to their relatively complex life cycle, as it is often the case in cooperative breeders (Hamilton and Taborsky 2005, Ozgul et al. 2014). Relatively more complex life cycles have been previously linked to demographic buffering (Koons et al. 2009). In the case of meerkats, the presence of subordinates, which help babysit the dominant pair’s offspring, may result in less variable recruitment than would otherwise be expected in the harsh environment meerkats inhabit (Clutton-Brock et al. 1998). However, recent evidence linking vital rates to climatic variables suggests that all vital rates in meerkats can substantially vary with climate depending on group and population structure (Paniw et al. 2019). These nuances are difficult to capture using simple correlation measures to assess demographic strategies.

Further evidence of how context dependence in population dynamics can produce variable variance-sensitivity relationships in vital rates comes from looking at the population growth rates of the study populations (see Table S2 on calculations of population growth rates). Populations with growth rates < 1 are decreasing in size, suggesting that they are living outside of their optimal environmental conditions (Wilmers et al. 2007, but see Csergő et al. 2017). We can expect these populations to experience unnatural fluctuations in their vital rates (Regehr et al. 2010). Indeed, populations in our dataset which were decreasing (Table S2) showed high variation in critical vital rates (Fig. S2). This was also true for the chimpanzee population, which we would expect to be buffered from environmental variation. A tendency towards lability in this and other species may therefore be an artifact of the population context.

Overall, the animal populations we studied represented a taxonomically biased sample of animal life histories, as data on temporal variation in vital rates for amphibians, fishes and insects, which have been shown to strongly track environmental variation, do not exist - or at least not readily open access (for amphibians: (Cayuela et al. 2017); for insects: (Valtonen et al. 2014); for fishes: (Schlosser 1990)). With larger sample sizes of less frequently studied taxa, we may expect stronger evidence for lability. Finally, in species that rely importantly on movement, such as migrating birds, proper estimates of buffering and lability can be harder to calculate, given the difficulty of measuring all relevant vital rates in all locations occupied by a population over time (Hostetler et al. 2015).

In conclusion, our study highlights that various factors such as the social structure of the populations, microhabitat conditions, and non-equilibrium population dynamics, can cause variation in vital rates that would not be expected under the theory of demographic buffering or lability. We emphasize that future comparative studies looking at vital-rate variations across taxa should explicitly link vital-rate responses to relevant drivers to better capture and account for context dependence.

## Data accessibility

Data and scripts for this publication are available in the Supporting Information and online at the Dryad Digital Repository: <placeholder doi>. The demographic data from most of the matrix population models used in this study are also archived at www.comadre-db.org. See Table S1 for a full list of citations of the original sources.

## Notes

### Competing Interest Statement

The authors have declared no competing interest.

## References

Adler, P. B. and Drake, J. M. 2008. Environmental variation, stochastic extinction, and competitive coexistence. - Am. Nat. 172: 186–195.

Bjørkvoll, E. et al. 2016. Demographic buffering of life histories? Implications of the choice of measurement scale. - Ecology 97: 40–47.

Bonnet, T. et al. 2017. Bigger is Fitter? Quantitative genetic decomposition of selection reveals an adaptive evolutionary decline of body mass in a wild rodent population. - PLoS Biol. 15: e1002592.

Boyce, M. et al. 2006. Demography in an increasingly variable world. - Trends Ecol. Evol. 21: 141–148.

Brooks, S. P. and Gelman, A. 1998. General methods for monitoring convergence of iterative simulations. - J. Comput. Graph. Stat. 7: 434–455.

Burns, J. H. et al. 2010. Empirical tests of life-history evolution theory using phylogenetic analysis of plant demography. - J. Ecol. 98: 334–344.

Campos, F. A. et al. 2017. Does climate variability influence the demography of wild primates? Evidence from long-term life-history data in seven species. - Glob. Chang. Biol. 23: 4907–4921.

Caswell, H. 2001. Matrix Population Models: Construction, Analysis, and Interpretation. - Sinauer Associates Incorporated.

Caswell, H. 2007. Sensitivity analysis of transient population dynamics. - Ecol. Lett. 10: 1–15.

Cayuela, H. et al. 2017. Life history tactics shape amphibians’ demographic responses to the North Atlantic Oscillation. - Glob. Chang. Biol. 23: 4620–4638.

Chevin, L.-M. et al. 2010. Adaptation, plasticity, and extinction in a changing environment: Towards a predictive theory. - PLoS Biology 8: e1000357.

Clutton-Brock, T. H. et al. 1998. Costs of cooperative behaviour in suricates (*Suricata suricatta*). - Proc. Biol. Sci. 265: 185–190.

Compagnoni, A. et al. 2016. The effect of demographic correlations on the stochastic population dynamics of perennial plants. - Ecol. Monogr. 86: 480–494.

Csergő, A. M. et al. 2017. Less favourable climates constrain demographic strategies in plants. - Ecol. Lett. 20: 969–980.

DeCesare, N. J. et al. 2012. Estimating ungulate recruitment and growth rates using age ratios. - J. Wildl. Manage. 76: 144–153.

de Kroon, H. et al. 1986. Elasticity: The relative contribution of demographic parameters to population growth rate. - Ecology 67: 1427–1431.

Doak, D. F. et al. 2005. Correctly estimating how environmental stochasticity influences fitness and population growth. - Am. Nat. 166: E14–21.

Drake, J. M. 2005. Population effects of increased climate variation. - Proc. Biol. Sci. 272: 1823–1827.

Drake, J. M. and Lodge, D. M. 2004. Effects of environmental variation on extinction and establishment. - Ecol. Lett. 7: 26–30.

Franco, M. and Silvertown, J. 2004. A comparative demography of plants based upon elasticities of vital rates. - Ecology 85: 531–538.

Gillespie, J. H. 1977. Natural selection for variances in offspring numbers: A new evolutionary principle. - Am. Nat. 111: 1010–1014.

Hadfield, J. D. 2010. MCMCglmm: MCMC methods for multi-response GLMMs in R. – J. Stat. Softw. 33: 1–22.

Hamilton, I. M. and Taborsky, M. 2005. Unrelated helpers will not fully compensate for costs imposed on breeders when they pay to stay. - Proc. Biol. Sci. 272: 445–454.

Healy, K. et al. 2019. Animal life history is shaped by the pace of life and the distribution of age-specific mortality and reproduction. – Nat. Ecol. Evol. 3: 1217–1224.

Hervé, M. 2019. RVAideMemoire: testing and plotting procedures for biostatistics. - R package version 0. 9--73 in press.

Hilde, C. H. et al. 2020. The demographic buffering hypothesis: Evidence and challenges. – Trend Ecol. Evol. 35: 523–538.

Hinchliff, C. E. et al. 2015. Synthesis of phylogeny and taxonomy into a comprehensive tree of life. - Proc. Natl. Acad. Sci. U. S. A. 112: 12764–12769.

Hostetler, J. A. et al. 2015. Full-annual-cycle population models for migratory birds. - Auk 132: 433–449.

Intergovernmental Panel on Climate Change 2014. Climate Change 2014 - Impacts, Adaptation and Vulnerability: Part B: Regional Aspects: Volume 2, Regional Aspects: Working Group II Contribution to the IPCC Fifth Assessment Report. - Cambridge University Press.

Jongejans, E. et al. 2010. Plant populations track rather than buffer climate fluctuations. - Ecol. Lett. 13: 736–743.

Koons, D. N. et al. 2009. Is life-history buffering or lability adaptive in stochastic environments? - Oikos 118: 972–980.

Lawson, C. R. et al. 2015. Environmental variation and population responses to global change. - Ecol. Lett. 18: 724–736.

Lewontin, R. C. and Cohen, D. 1969. On population growth in a randomly varying environment. - Proc. Natl. Acad. Sci. U. S. A. 62: 1056–1060.

Li, S.-L. and Ramula, S. 2015. Demographic strategies of plant invaders in temporally varying environments. - Popul. Ecol. 57: 373–380.

Link, W. A. and Doherty, P. F. 2002. Scaling in sensitivity analyses. - Ecology 83: 3299–3305.

McDonald, J. L. et al. 2017. Divergent demographic strategies of plants in variable environments. – Nat. Ecol. Evol. 1: 29.

Michonneau, F. et al. 2016. rotl: An R package to interact with the Open Tree of Life data. - Methods Ecol. Evol. 7: 1476–1481.

Miller, D. A. et al. 2011. Stochastic population dynamics in populations of western terrestrial garter snakes with divergent life histories. - Ecology 92: 1658–1671.

Moore, S. E. and Huntington, H. P. 2008. Arctic marine mammals and climate change: impacts and resilience. - Ecol. Appl. 18: s157–165.

Morris, W. F. and Doak, D. F. 2004. Buffering of life histories against environmental stochasticity: Accounting for a spurious correlation between the variabilities of vital rates and their contributions to fitness. - Am. Nat. 163: 579–590.

Morris, W. F. et al. 2008. Longevity can buffer plant and animal populations against changing climatic variability. - Ecology 89: 19–25.

Morris, W. F. et al. 2011. Low demographic variability in wild primate populations: fitness impacts of variation, covariation, and serial correlation in vital rates. - Am. Nat. 177: 14–28.

Myhrvold, N. P. et al. 2015. An amniote life-history database to perform comparative analyses with birds, mammals, and reptiles. - Ecology 96: 3109–3000.

Ozgul, A. et al. 2014. Linking body mass and group dynamics in an obligate cooperative breeder. - J. Anim. Ecol. 83: 1357–1366.

Paniw, M. et al. 2018. Interactive life-history traits predict sensitivity of plants and animals to temporal autocorrelation. - Ecol. Lett. 21: 275–286.

Paniw, M. et al. 2019. Life history responses of meerkats to seasonal changes in extreme environments. - Science 363: 631–635.

Paniw, M. et al. 2020. Assessing seasonal demographic covariation to understand environmental-change impacts on a hibernating mammal. - Ecol. Lett. 23: 588–597.

Parmesan, C. and Yohe, G. 2003. A globally coherent fingerprint of climate change impacts across natural systems. - Nature 421: 37–42.

Pfister, C. A. 1998. Patterns of variance in stage-structured populations: evolutionary predictions and ecological implications. - Proc. Natl. Acad. Sci. U. S. A. 95: 213–218.

Pusey, A. E. et al. 2008. Human impacts, disease risk, and population dynamics in the chimpanzees of Gombe National Park, Tanzania. - Am. J. Primatol. 70: 738–744.

Regehr, E. V. et al. 2010. Survival and breeding of polar bears in the southern Beaufort Sea in relation to sea ice. - J. Anim. Ecol. 79: 117–127.

Revell, L. J. 2012. phytools: An R package for phylogenetic comparative biology (and other things). - Methods Ecol. Evol. 3: 217–223.

Ripley, B. J. and Caswell, H. 2006. Recruitment variability and stochastic population growth of the soft-shell clam, *Mya arenaria*. - Ecol. Modell. 193: 517–530.

Rodríguez□Caro, R. C. et al. 2021. The limits of demographic buffering in coping with environmental variation. - Oikos in press.

Rotella, J. J. et al. 2012. Evaluating the demographic buffering hypothesis with vital rates estimated for Weddell seals from 30 years of mark-recapture data. - J. Anim. Ecol. 81: 162–173.

Salguero-Gómez, R. et al. 2016. COMADRE: a global data base of animal demography. - J. Anim. Ecol. 85: 371–384.

Schlosser, I. J. 1990. Environmental variation, life history attributes, and community structure in stream fishes: Implications for environmental management and assessment. - Environ. Manage. 14: 621–628.

Tuljapurkar, S. D. 1982. Population dynamics in variable environments. III. Evolutionary dynamics of r-selection. - Theor. Popul. Biol. 21: 141–165.

Valtonen, A. et al. 2014. Is climate warming more consequential towards poles? The phenology of Lepidoptera in Finland. - Glob. Chang. Biol. 20: 16–27.

van Tienderen, P. H. 1995. Life cycle trade-offs in matrix population models. - Ecology 76: 2482–2489.

Williams, J. M. et al. 2008. Causes of death in the Kasekela chimpanzees of Gombe National Park, Tanzania. - Am. J. Primatol. 70: 766–777.

Wilmers, C. C. et al. 2007. A perfect storm: the combined effects on population fluctuations of autocorrelated environmental noise, age structure, and density dependence. - Am. Nat. 169: 673–683.

Zúñiga-Vega, J. J. et al. 2007. Analysis of the population dynamics of an endangered lizard (*Xenosaurus grandis*) through the use of projection matrices. - Copeia 2007: 324–335.

