## Supplementary Materials for "Beyond demographic buffering: Context dependence in demographic strategies across animals"

**Data and *R* scripts**

All supporting data to run the *R* scripts, as well as data generated by the *R* scripts are available on Dryad and should be saved in one local folder. The setwd() function in the *R* scripts should be modified to link to this folder.

To replicate the analyses in this ms, the user is advised to run the *R* scripts in the following order:

1. corrected sensitivities and variances.R: Uses annual matrix population models from 31 animal populations to obtain vital rates, variance-stabilized sensitivities of the population growth rate to these vital rates, and standardized variation in the vital rates.
2. corrected integrated sensitivities and variances.R: Replicates analyses in (1), but calculates integrated sensitivities following van Tienderen (1995).
3. LH Traits.R: Calculates age at sexual maturity and the Gini index, and models these two traits as functions of body mass and matrix dimension, while controlling for phylogeny, using *MCMCglmm*. This script is based largely on code developed by Healy and co-authors (2019), and these authors should be cited in case this script is useful for readers.
4. MCMC_glmm.R: Models log(variance-stabilized sensitivities) of the population growth rate to vital rates as a function of the standardized variation in the vital rates, while accounting for the effect of life-history traits (residuals from models in [3]) on the slope of this relationship and controlling for phylogeny, using *MCMCglmm*. The structure of this script is based on code developed by Healy and co-authors (2019).
5. MCMC_glmm_integrated.R: As script in (4) but response is variance-stabilized integrated sensitivities of the population growth rate to vital rates.

**Supporting Materials S1.** Extra tables and details on the methods

**Table S1**. List of animal species used in this study. * citations refer to the original studies from which we obtained the MPMs in the COMADRE database.

| Species name | Common name | Class | Source |
| --- | --- | --- | --- |
| *Anser caerulescens* | Snow goose | Aves | (Saether and Bakke 2000, Cooch et al. 2001) |
| *Anthropoides paradiseus* | Blue crane | Aves | (Altwegg and Anderson 2009)* |
| *Callospermophilus lateralis* | Golden-mantled ground squirrel | Mammalia | (Hostetler et al. 2012)* |
| *Cebus capucinus* | White-headed capuchin | Mammalia | (Morris et al. 2011)* |
| *Centrocercus minimus* | Gunnison grouse | Aves | (Davis et al. 2014)* |
| *Cercopithecus mitis* | Blue monkey | Mammalia | (Morris et al. 2011)* |
| *Chionomys nivalis* | Snow vole | Mammalia | (Koons et al. 2009, Bonnet et al. 2017) |
| *Chlorocebus aethiops* | Grivet | Mammalia | (Isbell et al. 2009, Morris et al. 2011) |
| *Ciconia ciconia* | White stork | Aves | (Saether and Bakke 2000, Schaub et al. 2004) |
| *Erythrocebus patas* | Patas monkey | Mammalia | (Isbell et al. 2009, Morris et al. 2011) |
| *Falco naumanni* | Lesser kestrel | Aves | (Hiraldo et al. 1996)* |
| *Falco peregrinus* | Peregrine falcon | Aves | (Altwegg et al. 2013)* |
| *Gorilla beringei* | Eastern gorilla | Mammalia | (Morris et al. 2011)* |
| *Macaca mulatta* | Rhesus macaque | Mammalia | (Morris et al. 2011, Kessler et al. 2014) |
| *Notamacropus eugenii* | Tammar wallaby | Mammalia | (Chambers and Bencini 2010)* |
| *Marmota flaviventris* | Yellow-bellied marmot | Mammalia | (Paniw et al. 2020) |
| *Orcinus orca* | Killer whale | Mammalia | (Vélez-Espino et al. 2014)* |
| *Ovis aries* | Soay sheep | Mammalia | (Clutton-Brock et al. 1992)* |
| *Pan troglodytes schweinfurthii* | Eastern chimpanzee | Mammalia | (Morris et al. 2011)* |
| *Papio cynocephalus* | Yellow baboon | Mammalia | (Morris et al. 2011)* |
| *Propithecus verreauxi* | Verreaux’s sifaka | Mammalia | (Morris et al. 2011)* |
| *Pygoscelis adeliae* | Adélie penguin | Aves | (Hinke et al. 2017)* |
| *Rangifer tarandus caribou* | Woodland caribou | Mammalia | (DeCesare et al. 2012) |
| *Strix occidentalis occidentalis* | California spotted owl | Aves | (Saether and Bakke 2000, LaHaye et al. 2004) |
| *Suricata suricatta* | Meerkat | Mammalia | (Paniw et al. 2019) |
| *Turdus torquatus* | Ring ouzel | Aves | (Sim et al. 2010)* |
| *Umbonium costatum* | - | Gastropoda | (Noda and Nakao 1996)* |
| *Xenosaurus grandis* | Knob-scaled lizard | Reptilia | (Zúñiga-Vega et al. 2007)* |
| *Xenosaurus sp.* | - | Reptilia | (Zamora-Abrego et al. 2010)* |

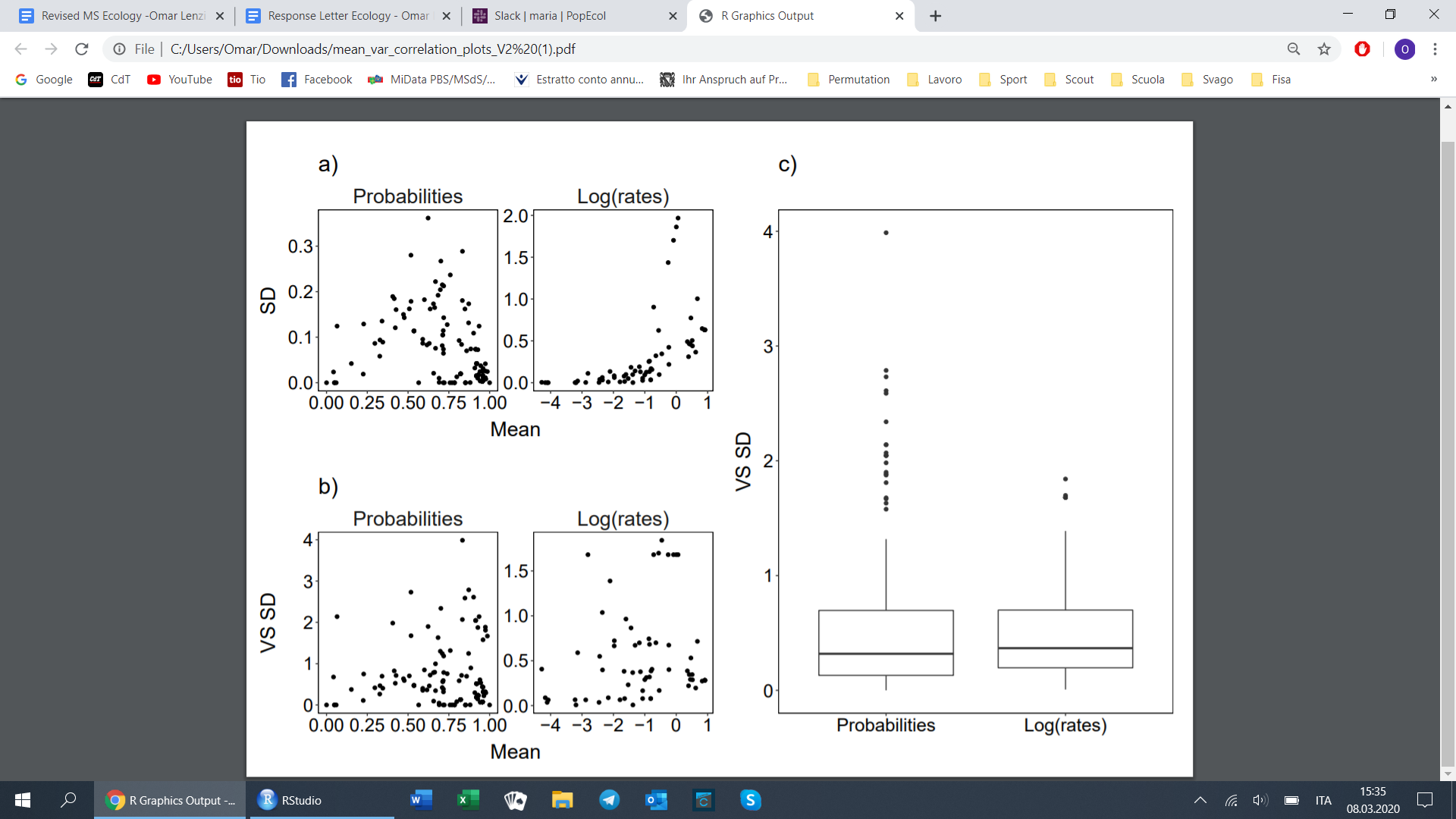

**Figure S1.** Relationship between the mean and standard deviation (SD) of (a) unscaled demographic probabilities and rates and (b) after variance-standardizing the standard deviation (VS SD). Logit-scaling demographic probabilities removed the constraints on the variance of probabilities while log-scaling demographic rates removed the pattern of increased variance with mean. c) The link-scaled measures of variance in probabilities and rates are spread over a similar distribution (Wilcoxon test W = 4994, p = 0.75) even though they are measured on different scales.

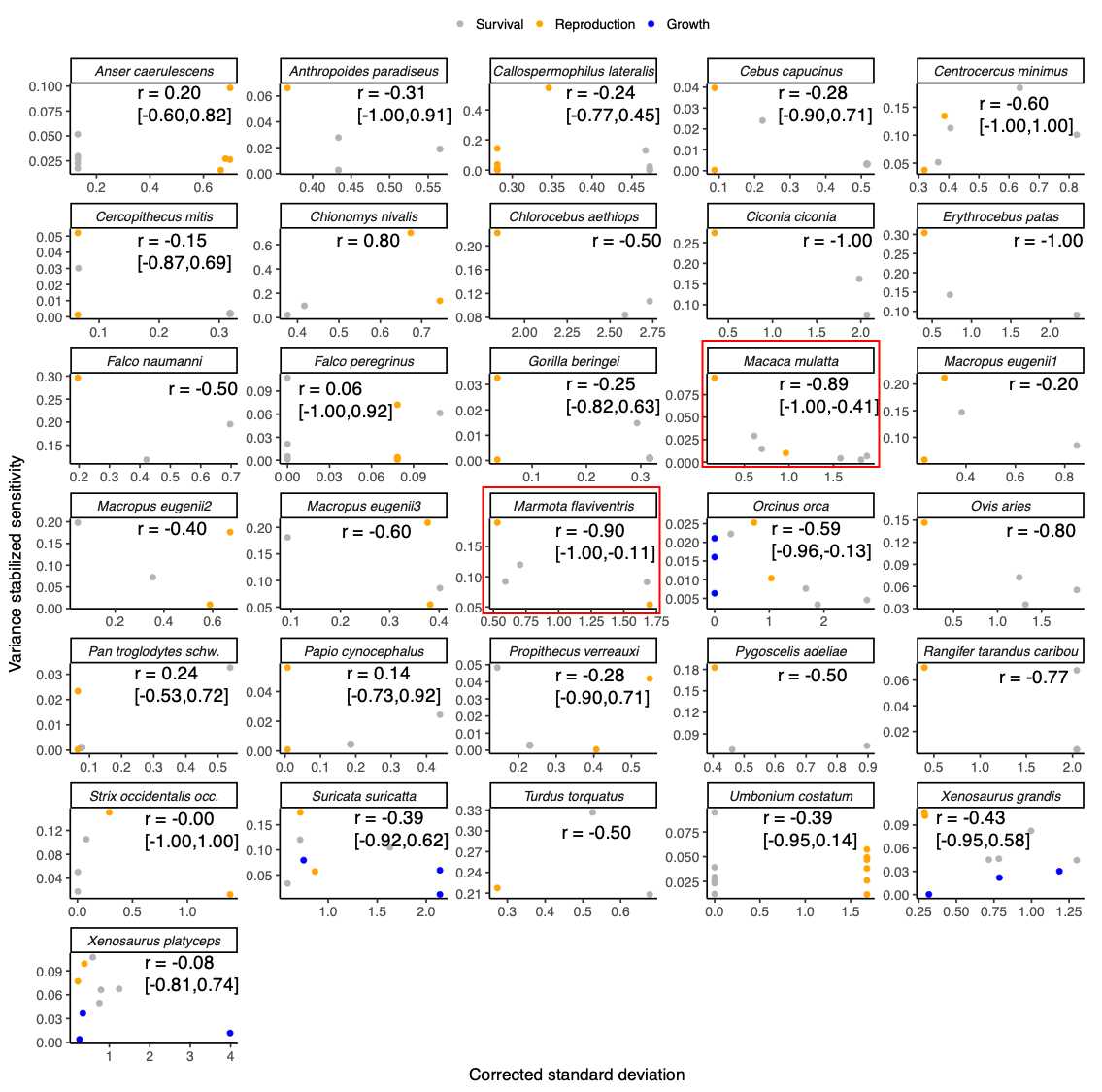

**Figure S2.** Temporal variation of stage-specific vital rates (*i.e.* the corrected standard deviation) and their importance to the population growth rate (*i.e.* the variance stabilized sensitivity) for each study population. *r* is the Spearman correlation coefficient of sensitivities and standard deviations. C.I. of *r* are indicated in brackets. The two species for which the C.I. does not cross 0 are framed in red.

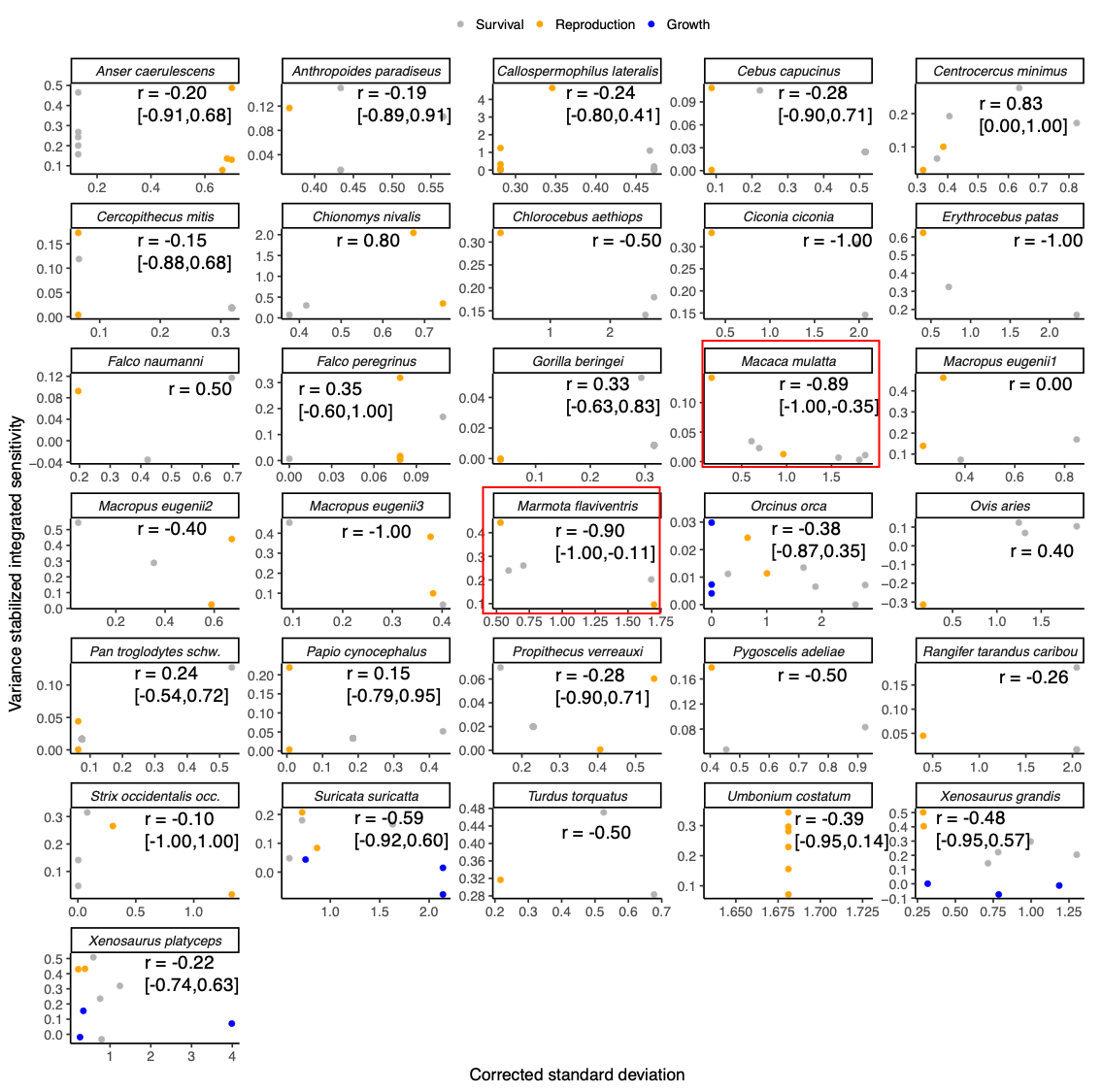

**Figure S3.** Temporal variation of stage-specific vital rates (*i.e.* the corrected standard deviation) and their importance to the population growth rate (*i.e.* the variance stabilized integrated sensitivity) for each study population. Integrated sensitivities were calculated following (van Tienderen 1995) and account for the covariation among vital rates. *r* is the Spearman correlation coefficient of sensitivities and standard deviations. C.I. of *r* are indicated in brackets. The two species for which the C.I. does not cross 0 are framed in red.

**Table S2.** Summary of the values of the four life-history traits and the stochastic population growth rate λ_s_ calculated for each of the 31 animal populations corresponding to 29 animal species in this study. *L_𝛼_*= Age at sexual maturity, G = Gini index, describing the spread of reproduction, with G = 0 indicating a fully semelparous and G = 1 indicating a extreme iteroparous organism. To distinguish multiple populations of the same species we used numbers 1-3 (*Macropus eugenii*). λs = stochastic population growth rate.

| Species name | *L_𝛼_* | *G* | *Mass (g)^#^* | λ_s_ |
| --- | --- | --- | --- | --- |
| *Anser caerulescens* | 2.00 | 0.68 | 2630 | 1.09 |
| *Anthropoides paradiseus* | 5.00 | 0.69 | 4341 | 1.05 |
| *Callospermophilus lateralis* | 1.00 | 0.63 | 300 | 1.97 |
| *Cebus capucinus* | 7.00 | 0.70 | 2655 | 1.02 |
| *Centrocercus minimus* | 3.00 | 0.72 | 1650 | 0.64 |
| *Cercopithecus mitis* | 8.00 | 0.72 | 6116 | 1.04 |
| *Chionomys nivalis* | 1.00 | 0.77 | 42 | 0.83 |
| *Chlorocebus aethiops* | 2.00 | 0.70 | 5104 | 1.15 |
| *Ciconia ciconia* | 2.00 | 0.69 | 3350 | 1.29 |
| *Erythrocebus patas* | 2.00 | 0.73 | 7660 | 1.09 |
| *Falco naumanni* | 2.00 | 0.69 | 152 | 1.22 |
| *Falco peregrinus* | 2.00 | 0.62 | 827 | 0.94 |
| *Gorilla beringei* | 10.00 | 0.71 | 139842 | 1.03 |
| *Macaca mulatta* | 4.00 | 0.71 | 6614 | 1.12 |
| *Marmota flaviventris* | 2.00 | 0.66 | 2800 | 1.10 |
| *Macropus eugenii 1* | 1.00 | 0.73 | 5037 | 1.00 |
| *Macropus eugenii 2* | 1.00 | 0.73 | 5037 | 0.88 |
| *Macropus eugenii 3* | 1.00 | 0.73 | 5037 | 0.91 |
| *Orcinus orca* | 8.91 | 0.64 | 3500000 | 1.00 |
| *Ovis aries* | 3.00 | 0.70 | 24000 | 1.08 |
| *Pan troglodytes schweinfurthii* | 16.00 | 0.72 | 44983 | 0.98 |
| *Papio cynocephalus* | 7.00 | 0.71 | 17851 | 1.05 |
| *Propithecus verreauxi* | 7.00 | 0.71 | 3588 | 0.99 |
| *Pygoscelis adeliae* | 2.00 | 0.7 | 4800 | 1.13 |
| *Rangifer tarandus caribou* | 2.00 | 0.72 | 100000 | 0.99 |
| *Strix occidentalis occidentalis* | 2.00 | 0.70 | 609 | 1.03 |
| *Suricata suricatta* | 2.00 | 0.60 | 776 | 1.03 |
| *Turdus torquatus* | 2.00 | 0.65 | 109 | 0.56 |
| *Umbonium costatum* | 1.00 | 0.66 | 2 | 1.08 |
| *Xenosaurus grandis* | 1.00 | 0.69 | 23 | 1.16 |
| *Xenosaurus sp.* | 1.00 | 0.67 | 19 | 1.02 |

### #We obtained average adult body mass from the Amniote database (Myhrvold et al. 2015) or the relevant articles in Table S1.

### **Stochastic population growth rate**

### We simulated λ_s_ for each population by projecting population dynamics over a period of 450'000 years after discarding 50,000 initial time steps of simulations to decouple λ_s_ from initial conditions (Tuljapurkar et al. 2003). At each time step, we randomly (i.i.d. simulation) sampled one MPM from all the MPMs belonging to a focal population. We obtained log λ_s_ by calculating the time-averaged cumulative growth. We used random MPM sampling as this is the most common procedure in stochastic population simulations (Paniw et al. 2018) and was used in all of the studies but one that performed stochastic simulations on our study populations (seven studies in total).

### **Details on phylogenetic tree construction**

As an outgroup to root the tree, we chose *Xenoturbella bocki,* as it is a closely related species of the clade including all of our study species, without belonging to it. In one case, we had three populations of the same species (the tammar wallaby, *Notamacropus eugenii)*. We added two populations as extra branches to one of the populations (chosen arbitrarily), labeling them as *Macropus eugenii b* and *Macropus eugenii c*. In one case, a specimen was not identified to species level: *Xenosaurus sp.* *(Zamora-Abrego et al. 2010)*. In the phylogenetic analysis, *Xenosaurus platyceps* was used. Indeed, *X. sp* is considered by Zamora-Abrego et al. (2010) either as a new species, or a remarkably divergent population of *X. platyceps.* We then computed branch length of the tree and transformed it into an ultrametric tree using the package *ape* *(Paradis et al. 2004)*.

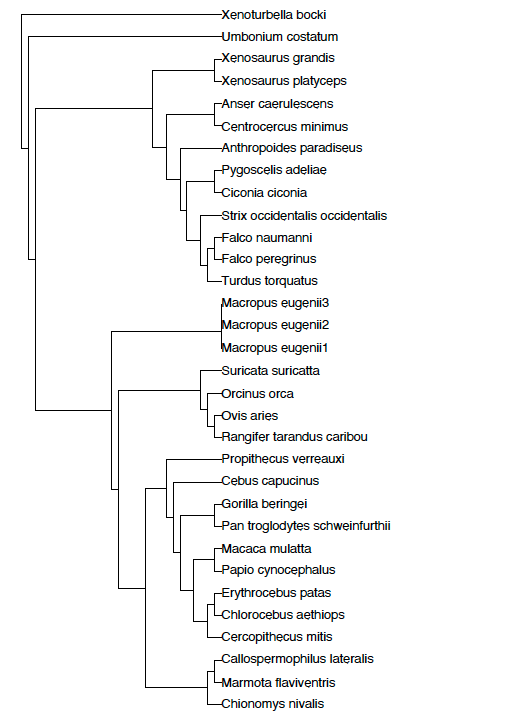

**Figure S4.** Phylogenetic tree of the 31 populations from 29 animal species for which we estimated demographic buffering and demographic lability. *Macropus eugenii* 1 - 3 represent three populations of *Macropus eugenii* (tammar wallaby). For the undefined species *Xenosaurus sp.* we used the most closely related species, *Xenosaurus platyceps. Xenoturbella bocki* represented the outgroup to root the tree.

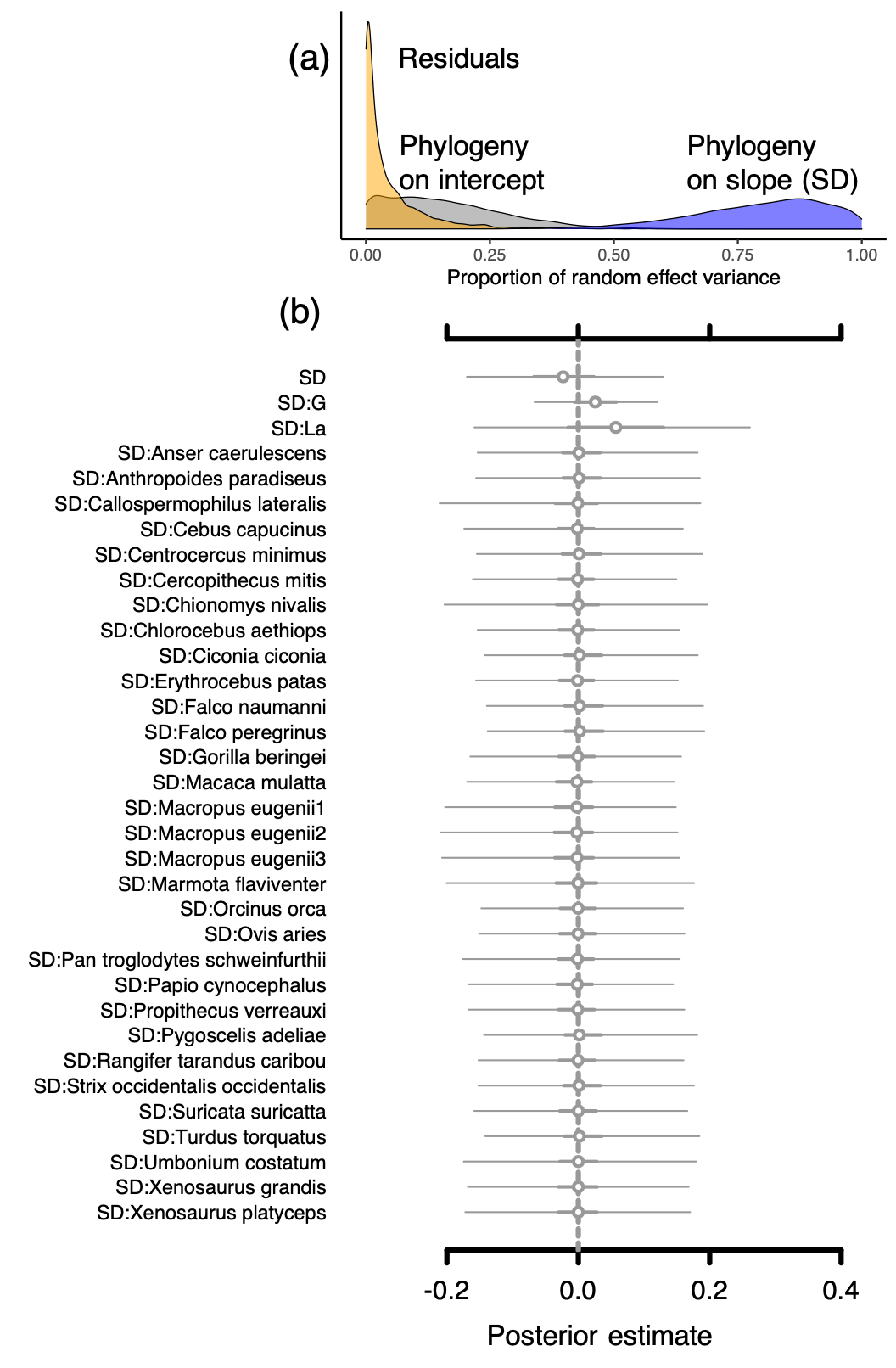

**Figure S5.** Effect of phylogeny (posterior parameter distribution) on model coefficients from the Bayesian mixed effect model describing the variance stabilized integrated sensitivity of population growth as a function of corrected vital rate standard deviations (SD) (a). Caterpillar plots (b) show the estimated fixed effects of SD, age at first reproduction (*L_𝛼_*) and reproduction spread (*G*), while accounting for a phylogenetically corrected species effect on SD. Interactions among predictors are depicted by “:”. Points represent posterior medians. Parameters where 50% credible intervals (C.I.) overlap 0 are indicated by open circles. Thick lines represent 50% C.I.; thin lines represent 95% credible intervals.

### **S1 References**

Altwegg, R. and Anderson, M. D. 2009. Rainfall in arid zones: possible effects of climate change on the population ecology of blue cranes. - Funct. Ecol. 23: 1014–1021.

Altwegg, R. et al. 2013. Nestboxes and immigration drive the growth of an urban Peregrine Falcon *Falco peregrinus* population. - Ibis 156: 107–115.

Bonnet, T. et al. 2017. Bigger is fitter? Quantitative genetic decomposition of selection reveals an adaptive evolutionary decline of body mass in a wild rodent population. - PLoS Biol. 15: e1002592.

Chambers, B. and Bencini, R. 2010. Road mortality reduces survival and population growth rates of tammar wallabies on Garden Island, Western Australia. - Wildl. Res. 37: 588.

Clutton-Brock, T. H. et al. 1992. Early Development and Population Fluctuations in Soay Sheep. - J. Anim. Ecol. 61: 381–396.

Cooch, E. et al. 2001. Retrospective analysis of demographic responses to environmental change: A lesser snow goose example. - Ecol. Monogr. 71: 377.

Davis, A. J. et al. 2014. An integrated modeling approach to estimating Gunnison sage-grouse population dynamics: combining index and demographic data. - Ecol. Evol. 4: 4247–4257.

DeCesare, N. J. et al. 2012. Estimating ungulate recruitment and growth rates using age ratios. - J. Wildl. Manage. 76: 144–153.

Healy, K. et al. 2019. Animal life history is shaped by the pace of life and the distribution of age-specific mortality and reproduction. – Nat. Ecol. Evol. 3: 1217–1224.

Hinke, J. T. et al. 2017. Variable vital rates and the risk of population declines in Adélie penguins from the Antarctic Peninsula region. - Ecosphere 8: e01666.

Hiraldo, F. et al. 1996. A demographic model for a population of the endangered lesser kestrel in southern Spain. - J. Appl. Ecol. 33: 1085.

Hostetler, J. A. et al. 2012. Stochastic population dynamics of a montane ground-dwelling squirrel. - PLoS One 7: e34379.

Isbell, L. A. et al. 2009. Demography and life histories of sympatric patas monkeys, *Erythrocebus patas*, and vervets, *Cercopithecus aethiops*, in Laikipia, Kenya. - Int. J. Primatol. 30: 103–124.

Kessler, M. J. et al. 2014. Long-term effects of tetanus toxoid inoculation on the demography and life expectancy of the Cayo Santiago rhesus macaques. - Am. J. Primatol. 77: 211–221.

Koons, D. N. et al. 2009. Is life-history buffering or lability adaptive in stochastic environments? - Oikos 118: 972–980.

LaHaye, W. S. et al. 2004. Temporal variation in the vital rates of an insular population of spotted owls (*Strix occidentalis occidentalis*): Contrasting effects of weather. - Auk 121: 1056.

Morris, W. F. et al. 2011. Low demographic variability in wild primate populations: fitness impacts of variation, covariation, and serial correlation in vital rates. - Am. Nat. 177: E14–28.

Myhrvold, N. P. et al. 2015. An amniote life-history database to perform comparative analyses with birds, mammals, and reptiles. - Ecology 96: 3109–3000.

Noda, T. and Nakao, S. 1996. Dynamics of an entire population of the subtidal snail *Umbonium costatum*: The importance of annual recruitment fluctuation. - J. Anim. Ecol. 65: 196–204.

Paniw, M. et al. 2018. Interactive life-history traits predict sensitivity of plants and animals to temporal autocorrelation. - Ecol. Lett. 21: 275–286.

Paniw, M. et al. 2019. Life history responses of meerkats to seasonal changes in extreme environments. - Science 363: 631–635.

Paniw, M. et al. 2020. Assessing seasonal demographic covariation to understand environmental-change impacts on a hibernating mammal. - Ecol. Lett. 23: 588–597.

Paradis, E. et al. 2004. APE: Analyses of Phylogenetics and Evolution in R language. - Bioinformatics 20: 289–290.

Saether, B.-E. and Bakke, O. 2000. Avian Life History Variation and Contribution of Demographic Traits to the Population Growth Rate. - Ecology 81: 642.

Schaub, M. et al. 2004. Is the reintroduced white stork (*Ciconia ciconia*) population in Switzerland self-sustainable? - Biol. Conserv. 119: 105–114.

Sim, I. M. W. et al. 2010. Characterizing demographic variation and contributions to population growth rate in a declining population. - J. Anim. Ecol. 80: 159–170.

Tuljapurkar, S. et al. 2003. The many growth rates and elasticities of populations in random environments. - Am. Nat. 162: 489–502.

van Tienderen, P. H. 1995. Life cycle trade-offs in matrix population models. - Ecology 76: 2482–2489.

Vélez-Espino, L. A. et al. 2014. Comparative demography and viability of northeastern Pacific resident killer whale populations at risk. - Canadian Technical Report of Fisheries and Aquatic Sciences 3084.

Zamora-Abrego, J. G. et al. 2010. Demography of a knob-scaled Lizard in Northeastern Querétaro, México. - Herpetologica 66: 39–51.

Zúñiga-Vega, J. J. et al. 2007. Analysis of the population dynamics of an endangered lizard (*Xenosaurus grandis*) through the use of projection matrices. - Copeia 2007: 324–335.
